## Supplemental Figures for "Protein Phosphatase 1 activity controls a balance between collective and single cell modes of migration"

### Supplemental Information

#### **Figure S1**, Related to Figure 1

(**A**) Fertility of control versus NiPp1-expressing females. The average progeny per female in each vial (individual plot points) is shown as a box-and-whiskers plot (see Figure 1 legend for details of plot). (**B**) Expression pattern of *c306*-GAL4 in the border cell cluster, as visualized by driving the expression of the membrane marker UAS-PLCδPH-GFP (green). Nuclei are labeled by DAPI (blue). The central polar cells (asterisk) express GFP. (**C**) Schematic drawing of the border cell cluster showing the patterns of GAL4 drivers. *upd*-GAL4 (green) is used to drive expression in polar cells; *slbo*-GAL4 (blue) is used to drive expression in outer border cells but not polar cells. (**D-G**) Overexpression of NiPp1 in border cells, driven by *slbo*-GAL4, disrupts border cell cluster migration and cohesion. (**D, E**) Stage 10 *slbo*-GAL4 egg chambers expressing mCD8-GFP (green), which is detected in border cells (arrowheads), and stained for DAPI to label nuclei (blue) and phalloidin to label F-actin (red, D). Control border cells (D) reach the oocyte as a single unit, but NiPp1 overexpressing border cells (E) dissociate from the cluster and fail to reach the oocyte. (**F**) Quantification of border cell migration for matched control and NiPp1 overexpression, shown as the percentage that did not complete (red), or completed (green) their migration to the oocyte (see Figure 1I for egg chamber schematic). (**G**) Quantification of cluster cohesion, shown as the percentage of border cells found as a single unit (1 part) or split into multiple parts (2-3 parts or >3 parts) in control versus NiPp1-expressing egg chambers. (**F, G**) ****$p < 0.0001$; unpaired two-tailed *t* test. Error bars represent SEM in 3 experiments, each trial assayed $n \geq 62$ egg chambers (total $n \geq 201$ for each genotype). (**H-K**) NiPp1 overexpression in polar cells, driven by *upd*-GAL4, does not impair border cell cluster migration or cohesion. (**H, I**) Stage 10 *upd*-GAL4 egg chambers expressing mCD8-ChRFP (red in H) or NiPp1-HA (red in I), which are detected in the polar cells (arrowheads), and co-stained for phalloidin to detect F-actin (magenta), SN to detect border cells (green), and DAPI to label nuclei (blue). (**J, K**) Quantification of border cell migration (J) and border cell cluster cohesion (K) for matched control and NiPp1overexpression. Error bars represent SEM in 3 experiments, each trial assayed $n \geq 80$ egg chambers (total $n \geq 313$ for each genotype). ****$p < 0.0001$, unpaired two-tailed t test. (**L-N**) Border cells expressing NiPp1 driven by *c306*-GAL4 (M, N), can separate from the polar cells, whereas control border cells (L) stay attached to polar cells. Stage 10 egg chambers stained for SN (green) to detect border cells, FasIII (red) to detect polar cells (arrows), and DAPI to label nuclei (blue). Yellow dashed line indicates anterior border of the oocyte. Example of a control (L) and two representative NiPp1 overexpressing border cell clusters (M, N), polar cells are indicated by magenta arrows. All genotypes are listed in Supplemental Table 2.

#### **Figure S2**, Related to Figure 1

(**A-B'**) Stage 10 wild-type (A, A') and NiPp1-expressing (B, B') egg chambers stained with anti-Slbo (green in A, B; white in A', B'), anti-Eya (red in A, B), and DAPI to detect nuclei (blue in A, B). Eya primarily marks the anterior follicle cells with lower levels in border cells but is absent from polar cells. Slbo marks border cells and polar cells (arrowheads). Insets, zoomed-in images of Slbo-expressing cells. (**C**) Quantification of cell number per cluster in control and

NiPp1-expressing border cell clusters. The total number of egg chambers scored for cell number in each genotype is shown and was assayed in 3 independent trials. Error bars represent SEM, *$p < 0.05$, unpaired two-tailed *t* test. All genotypes are listed in Supplemental Table 2.

**Figure S3**, Related to Figure 2

(**A-E**) Overexpression of Pp1c subunits on their own does not impair border cell migration to the oocyte. Stage 10 egg chambers of the indicated genotypes stained for Armadillo (Arm; β-Catenin) to detect cell membranes (green in A-D) or SN to detect border cells (green in E), HA to detect Pp1c overexpression (red) and DAPI to label nuclei (blue). (**F-K**) Overexpression of Pp1c subunits can rescue NiPp1-induced border cell migration defects and cohesion. Stage 10 egg chambers of the indicated genotypes stained for SN to detect border cells (green in G-K) or phalloidin to detect F-actin (green in F, red in G-K), mCD8-ChRFP (red in F), and DAPI to label nuclei (blue). Border cells in all panels are indicated by arrowheads. All genotypes are listed in Supplemental Table 2.

**Figure S4**, Related to Figures 2 and 3

(**A-B"**) NiPp1-HA overexpression promotes the nuclear localization of two Pp1c subunits, of Pp1α-96A and Flw. Stage 10 egg chambers co-expressing Pp1α-96A-GFP (green in A, A") or Flw-YFP (green in B, B") with UAS-NiPp1 were stained for anti-HA to detect NiPp1 expression (red in A, A', B, B') and DAPI to detect nuclei (blue in A, B; white in A', A", B', B"). Insets, zoomed-in images of border cells. (**C-C"**) Example of a stage 10 egg chamber with a *flw*$^{FP41}$ mutant clone, marked by the loss of nuclear mRFP (red in C, C'; dotted outline) and stained for SN (green in C") to mark border cells (arrowheads) and DAPI (blue in C) to mark nuclei. All genotypes are listed in Supplemental Table 2.

**Figure S5**, Related to Figure 4

(**A-F'**) Efficiency of cadherin-catenin *RNAi* in border cells as detected by antibody staining to the respective proteins. Stage 10 control (A, A', C, C', E, E'), *E-Cad-RNAi* (B, B'), *β-Cat-RNAi* (D, D'), and *α-Cat-RNAi* (F, F') egg chambers stained for SN (green), the respective proteins in red (E-Cad in A-B', β-Cat in C-D', and α-Cat in E-F'), and DAPI to label nuclei (blue in A, B, C, D, E, F). Border cells are indicated by arrowheads. Insets, zoomed-in views of border cell clusters. (**G, H**) Border cell migration (G) and cluster cohesion (H) in control versus *α-Cat-RNAi* driven by *upd*-GAL4 in the polar cells. Quantification at stage 10, $n \geq 44$ (total $n \geq 192$ for each genotype); ns, not significant, **$p < 0.01$, ***$p < 0.001$, unpaired two-tailed *t* test. (**G**) Quantification of migration shown as the percentage of egg chambers with complete (green), partial (blue), or no (red), border cell migration. (**H**) Quantification of cluster cohesion, shown as the percentage of border cells found as a single unit (no split, blue) or split into two or more parts (split, yellow). All genotypes are listed in Supplemental Table 2.

**Figure S6,** Related to Figure 5

(**A**) Close-up view of a live border cell cluster depicting how protrusions are measured. The main body of the border cell cluster is outlined (yellow circle). The protrusion length and area (green outline) were defined and measured as a cellular projection extending away from the main cluster or border cell. The schematic indicates how protrusion direction is defined. (**B, C**)

Quantification of protrusion max_length (B) and max_area (C) in control versus *Pp1c-RNAi* border cells. Data are presented as a box-and-whiskers plot (see Figure 1 legend for details of plot). *p < 0.05; **p < 0.01; ***p < 0.001; ****p < 0.0001; unpaired two-tailed *t* test. (**D, E**) Rac-FRET in wild-type versus NiPp1 border cells (n = 22 for control, n=18 for *slbo*>NiPp1). (D) Representative FRET images of control and NiPp1-expressing border cells, color-coded according to the heat map and FRET index. (E) Quantification of the total FRET index measured in control and NiPp1 border cells. Error bars represent SEM. p=0.0033 (**), unpaired *t* test with Welch's correction. All genotypes are listed in Supplemental Table 2.

**Figure S7,** Related to Figure 6

(**A-B'''''**) Pp1 restricts Myo-II to the cluster periphery. Stills from representative confocal videos of dynamic Sqh:GFP in mid-migration borders cells over the course of 5 minutes. (A-A''''') Control border cells (Video 15) have dynamic Sqh:GFP, which is mainly restricted to the cluster perimeter. Some signal is found in the central polar cells. (B-B''''') NiPp1 overexpressing border cells (Video 16) alters the localization of Sqh:GFP, with more uniform Sqh:GFP, especially at contacts between some border cells. All genotypes are listed in Supplemental Table 2.

**Figure S8,** Related to Figure7

(**A-J)** Mbs RNA (A-C) and protein (D-I") are found in border cells throughout migration (arrowheads, A-F). (**A-C**) *Mbs* RNA pattern as detected by *in situ* hybridization (green) in stage 9 to 10 egg chambers. Nuclear DNA, stained for DAPI, is shown in magenta. Images are from the Dresden Ovary Table http://tomancak-srv1.mpi-cbg.de/DOT/main[92]. (**D-F**) Wild-type (*w*$^{1118}$) egg chambers stained for Mbs protein (green) and DAPI to label nuclei (blue). (**G-J**) Colocalization of Mbs with Pp1c subunits in border cells. The border cells are co-stained for Mbs (red in G, G", I, I") and Pp1α-96A-GFP (green in G, G') or Flw-YFP (green in I, I'). DAPI labels the nuclei (blue). (H, J) Plot profiles of the fluorescent image intensities of Pp1α-96A-GFP (H), Flw-YFP (J), Mbs, and DAPI across the lines shown (G-G", I-I"). (**K-L''**) Mbs RNAi results in significant reduction of Mbs protein levels in border cells (L-L") compared to control (K-K"). Stage 10 egg chambers stained for Mbs (green in K, L; white in K', L') and DAPI to label nuclei (blue in K, L; white in K", L"). All genotypes are listed in Supplemental Table 2.

**Figure S9,** Related to Figure 7

(**A-C**) Pp1 promotes moderate levels of RhoA activity in border cells. (**A-B'**) Representative processed Rho-FRET images in control (A, A') and NiPp1 overexpressing (B, B') border cells. The CFP channel (A, B) is shown. The FRET images (A', B') are color-coded by the heat map. (**C**) Measurement of the total FRET index in matched control and NiPp1 overexpressing border cells. The total number of border cell clusters assayed is indicated. All genotypes are listed in Supplemental Table 2.

### Video Legends

**Video S1.** Control (*c306*-GAL4/+;*UAS*-mCherry::Jupiter/+) egg chamber showing normal border cell migration. Frames were acquired every 3 min with a 20x objective. Anterior is to the left.

**Video S2.** NiPp1 overexpressing (*c306*-GAL4/+;*UAS*-mCherry::Jupiter/+;*UAS*-NiPp1/+) egg chamber showing the migration defect and splitting phenotype. Frames were acquired every 3 min with a 20x objective. Anterior is to the left.

**Video S3.** Representative time-lapse video of a stage 9 NiPp1 overexpressing (*c306*-GAL4,tsGAL80/+;*UAS*-mCherry::Jupiter/+;*UAS*-NiPp1/+) egg chamber with DIC channel. Frames were acquired every 2 min with a 20x objective. Time is in hr:min. Anterior is to the left.

**Video S4.** Representative time-lapse video of a stage 9 NiPp1 overexpressing (*c306*-GAL4,tsGAL80/+;*UAS*- mCherry::Jupiter /+;*UAS*-NiPp1/+) egg chamber with DIC channel. Frames were acquired every 2 min with a 20x objective. Time is in hr:min. Anterior is to the left.

**Video S5.** NiPp1 overexpressing (*slbo*-GAL4/+; UAS-PLCδPH:GFP/UAS-NiPp1) egg chamber showing the loss of a membrane attachment between one border cell and the rest of the border cell cluster. Anterior is to the left.

**Video S6.** Control (*c306*-GAL4,tsGAL80/+; UAS-PLCδPH:GFP/+) egg chamber showing normal border cell migration. Frames were acquired every 3 min with a 20x objective. Anterior is to the left.

**Video S7.** Representative time-lapse video of a stage 9 *Pp1α-96A* RNAi (*c306*-GAL4,tsGAL80/+; *v27673*/+;UAS-PLCδPH:GFP/+) egg chamber. Frames were acquired every 3 min with a 20x objective. Anterior is to the left.

**Video S8.** Representative time-lapse video of a stage 9 *Pp1-87B* RNAi (*c306*-GAL4,tsGAL80/+; *v35024*/+;UAS-PLCδPH:GFP/+) egg chamber. Frames were acquired every 3 min with a 20x objective. Anterior is to the left.

**Video S9.** Representative time-lapse video of a stage 9 *Pp1-13C* RNAi (*c306*-GAL4,tsGAL80/+;*v*29058/+;UAS-PLCδPH:GFP/+) egg chamber. Frames were acquired every 3 min with a 20x objective. Anterior is to the left.

**Video S10.** Representative time-lapse video of a stage 9 *α-Catenin* RNAi (*c306*-GAL4,tsGAL80/+;*v107298/+;*UAS-PLCδPH:GFP/+) egg chamber. Frames were acquired every 3 min with a 20x objective. Anterior is to the left.

**Video S11.** Control (LifeAct-GFP/+) egg chamber showing the dynamics of F-actin with LifeAct-GFP, Frames were acquired every 2 min with a 40x water immersion objective. Anterior is to the left.

**Video S12.** NiPp1 overexpressing (*slbo*-Gal4/+;UAS-NiPp1/LifeAct-GFP) egg chamber showing F-actin dynamics with LifeAct-GFP. Frames were acquired every 2 min with a 40x water immersion objective. Anterior is to the left.

**Video S13.** Control (Sqh::GFP/+) egg chamber showing normal Sqh::GFP dynamics in early migration. Frames were acquired every 1 min with a 40x water immersion objective. Anterior is to the left.

**Video S14.** Representative NiPp1 overexpressing (*slbo*-Gal4/+;UAS-NiPp1/Sqh::GFP) egg Chamber showing the Sqh::GFP dynamics in early migration. Frames were acquired every 1 min with a 40x water immersion objective. Anterior is to the left.

**Video S15.** Control (Sqh::GFP/+) egg chamber showing normal Sqh::GFP dynamics in mid-migration. Frames were acquired every 1 min with a 40x water immersion objective. Anterior is to the left.

**Video S16.** Representative NiPp1 overexpressing (*slbo*-Gal4/+;UAS-NiPp1/Sqh::GFP) egg chamber showing the Sqh::GFP dynamics in mid-migration. Frames were acquired every 1 min with a 40x water immersion objective. Anterior is to the left.

Fig S1

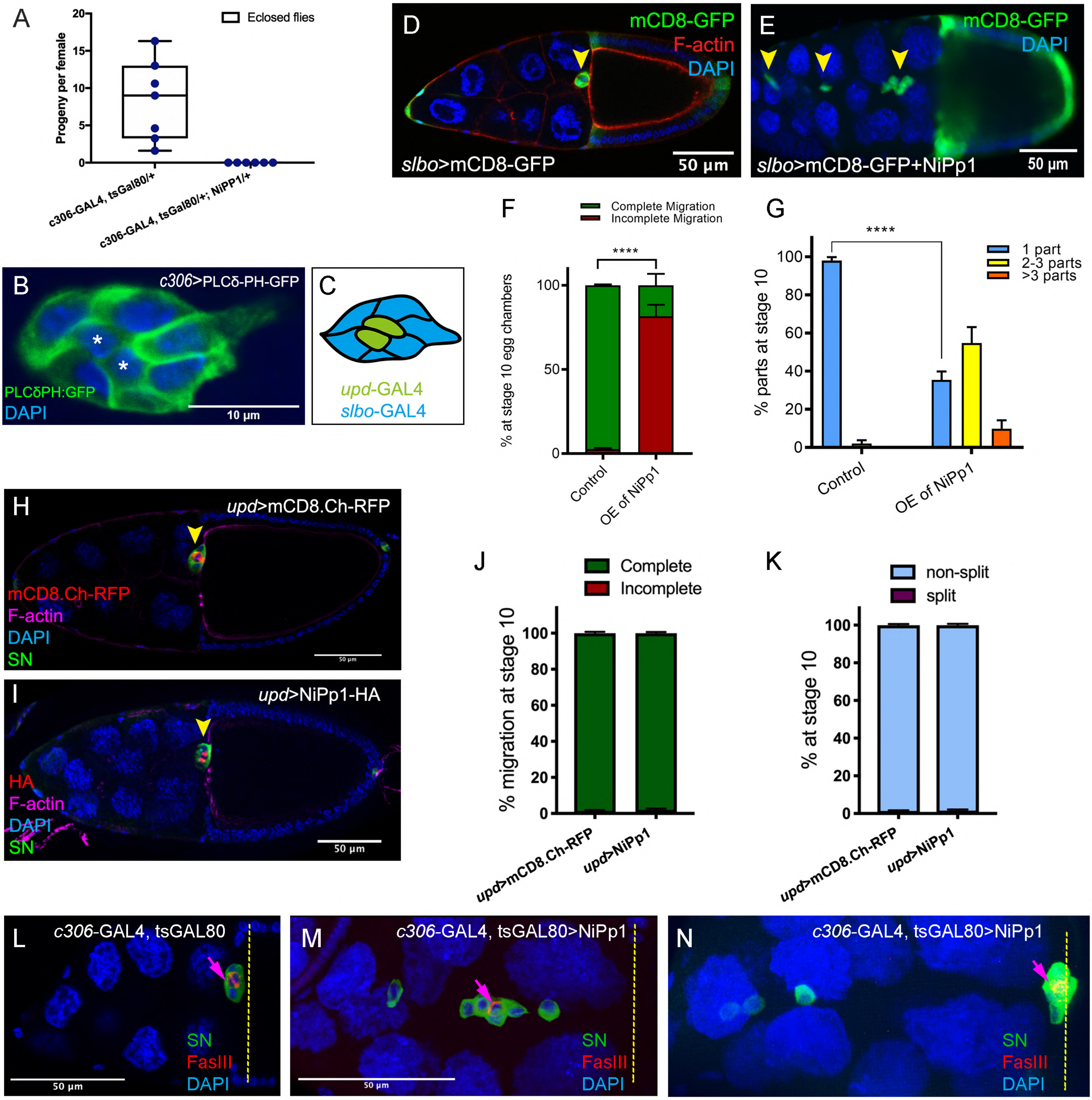

A
Eclosed flies
Progeny per female
c306-GAL4, tsGal80/+
c306-GAL4, tsGal80/+; NiPP1/+
B
c306>PLCδ-PH-GFP
PLCδPH:GFP
DAPI
10 μm
C
upd-GAL4
slbo-GAL4
D
mCD8-GFP
F-actin
DAPI
slbo>mCD8-GFP
50 μm
E
mCD8-GFP
DAPI
slbo>mCD8-GFP+NiPp1
50 μm
F
Complete Migration
Incomplete Migration
% at stage 10 egg chambers
****
Control
OE of NiPp1
G
1 part
2-3 parts
>3 parts
% parts at stage 10
****
Control
OE of NiPp1
H
upd>mCD8.Ch-RFP
mCD8.Ch-RFP
F-actin
DAPI
SN
I
upd>NiPp1-HA
HA
F-actin
DAPI
SN
50 μm
J
Complete
Incomplete
% migration at stage 10
upd>mCD8.Ch-RFP
upd>NiPp1
K
non-split
split
% at stage 10
upd>mCD8.Ch-RFP
upd>NiPp1
L
c306-GAL4, tsGAL80
SN
FasIII
DAPI
50 μm
M
c306-GAL4, tsGAL80>NiPp1
SN
FasIII
DAPI
50 μm
N
c306-GAL4, tsGAL80>NiPp1
SN
FasIII
DAPI

Fig S2

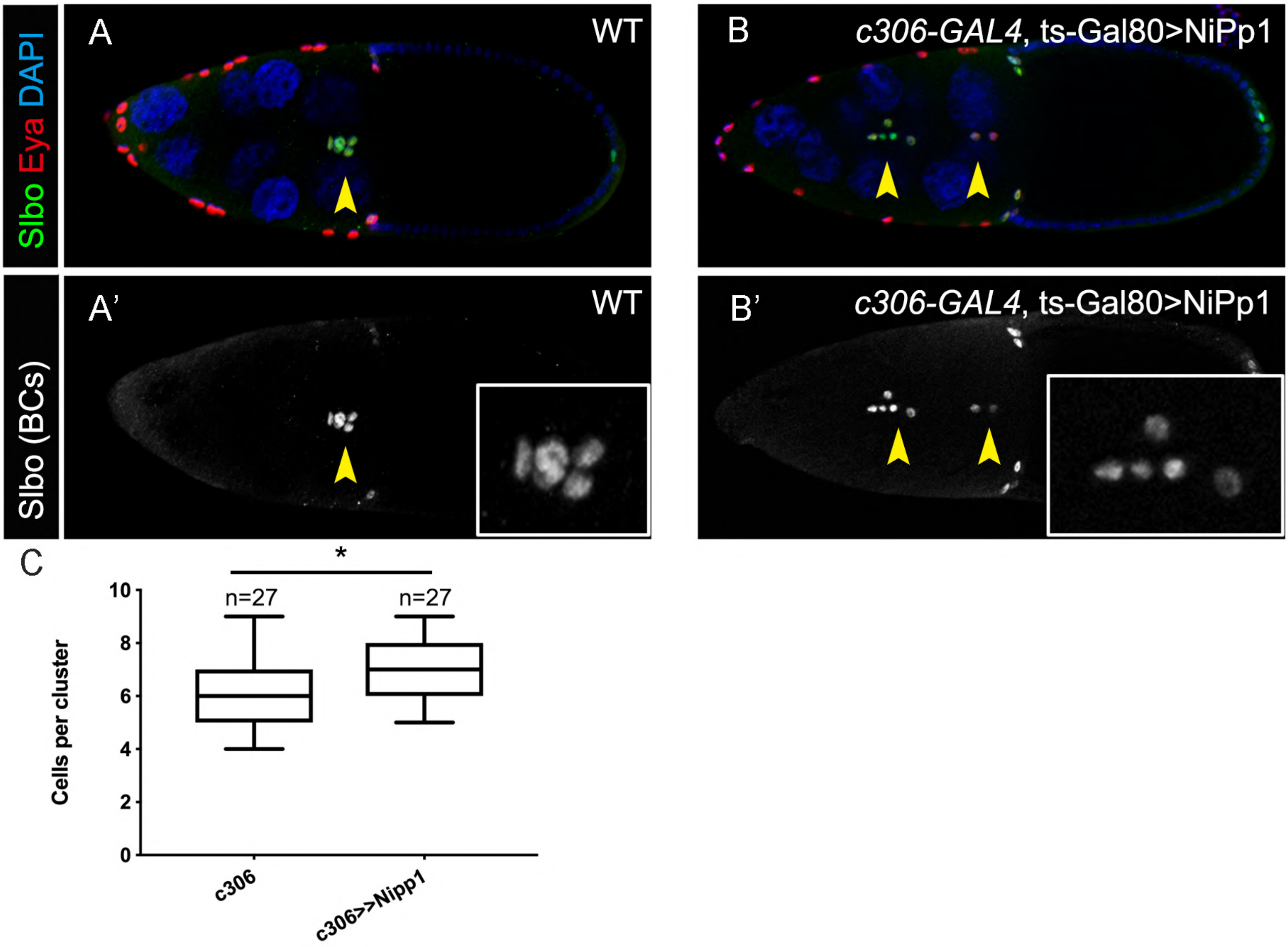

Slbo Eya DAPI
A
WT
B
c306-GAL4, ts-Gal80>NiPp1
Slbo (BCs)
A'
WT
B'
c306-GAL4, ts-Gal80>NiPp1
C
*
n=27
n=27
Cells per cluster
10
8
6
4
2
0
c306
c306>Nipp1

Fig S3

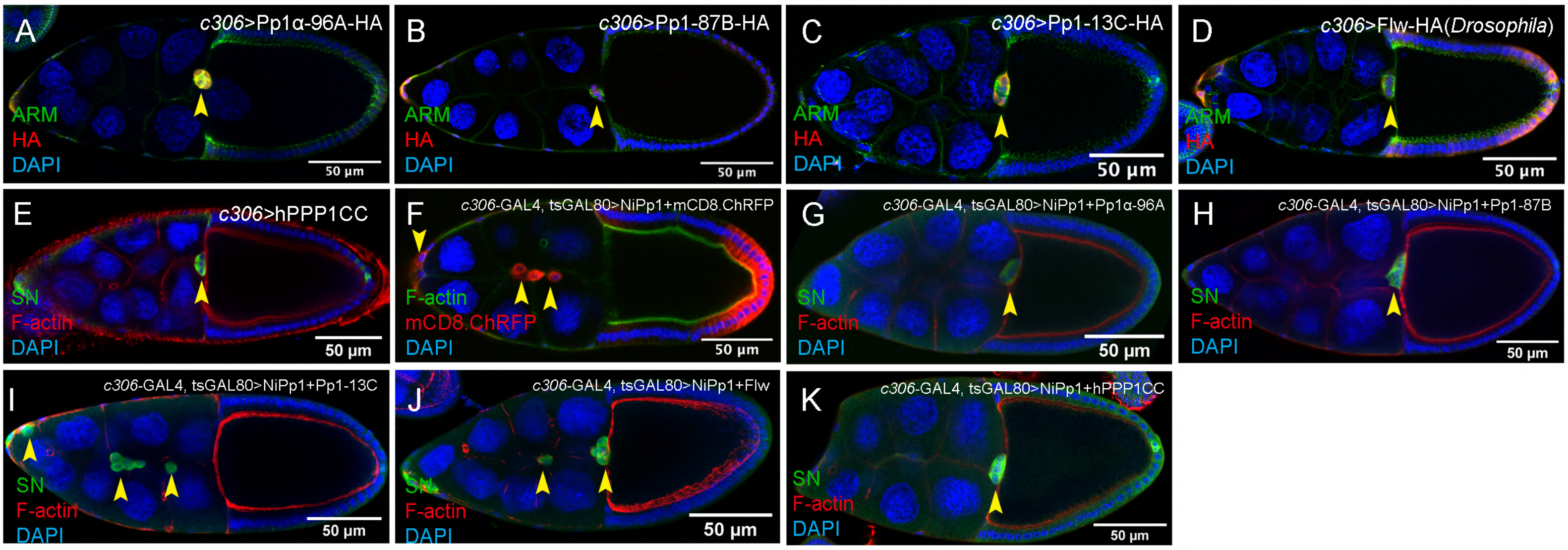

A
c306>Pp1α-96A-HA
ARM
HA
DAPI
50 µm
B
c306>Pp1-87B-HA
ARM
HA
DAPI
50 µm
C
c306>Pp1-13C-HA
ARM
HA
DAPI
50 µm
D
c306>Flw-HA(Drosophila)
ARM
HA
DAPI
50 µm
E
c306>hPPP1CC
SN
F-actin
DAPI
50 µm
F
c306-GAL4, tsGAL80>NiPp1+mCD8.ChRFP
F-actin
mCD8.ChRFP
DAPI
50 µm
G
c306-GAL4, tsGAL80>NiPp1+Pp1α-96A
SN
F-actin
DAPI
50 µm
H
c306-GAL4, tsGAL80>NiPp1+Pp1-87B
SN
F-actin
DAPI
50 µm
I
c306-GAL4, tsGAL80>NiPp1+Pp1-13C
SN
F-actin
DAPI
50 µm
J
c306-GAL4, tsGAL80>NiPp1+Flw
SN
F-actin
DAPI
50 µm
K
c306-GAL4, tsGAL80>NiPp1+hPPP1CC
SN
F-actin
DAPI
50 µm

Fig S4

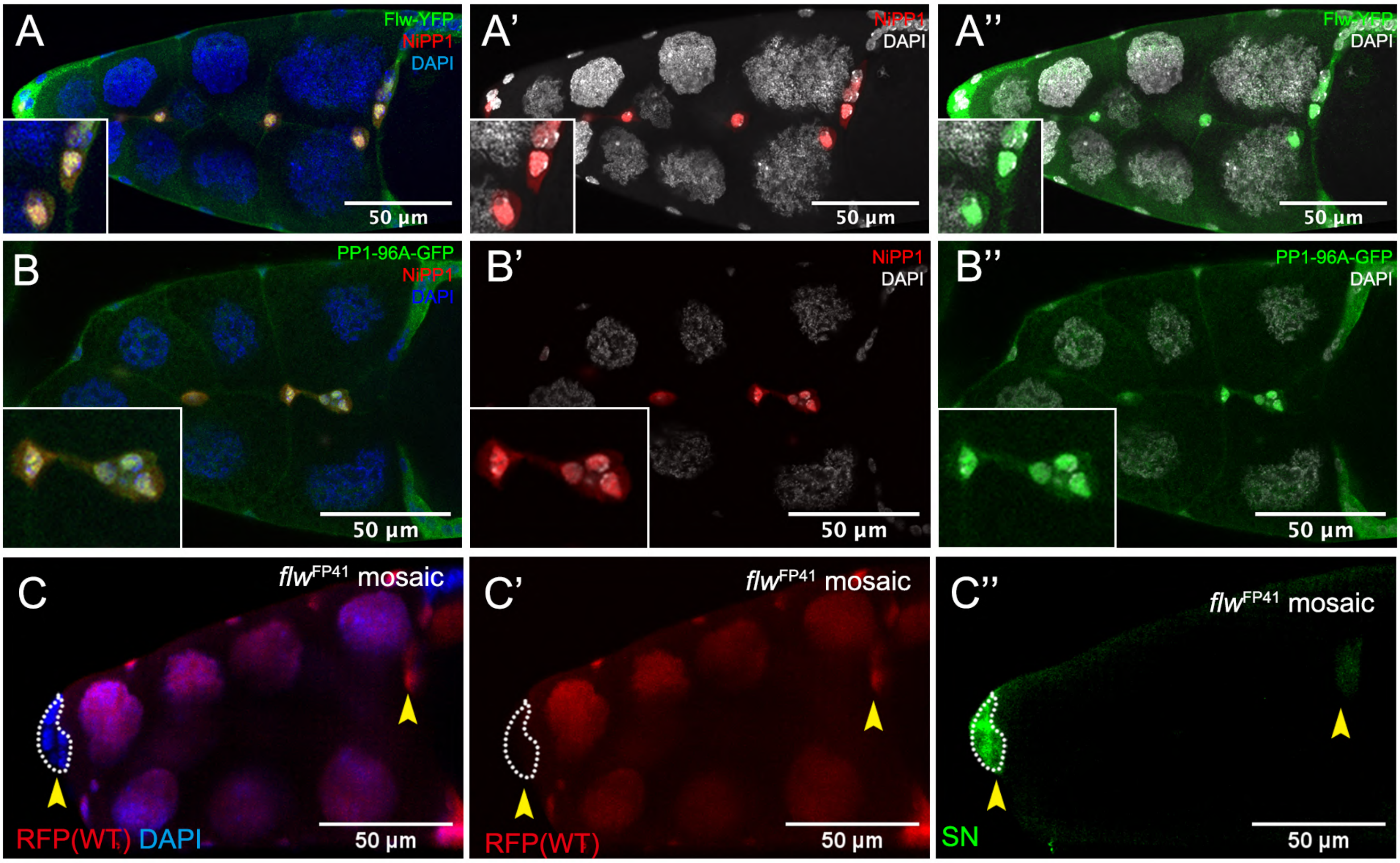

A
Flw-YFP
NiPP1
DAPI
50 µm
A'
NiPP1
DAPI
50 µm
A''
Flw-YFP
DAPI
50 µm
B
PP1-96A-GFP
NiPP1
DAPI
50 µm
B'
NiPP1
DAPI
50 µm
B''
PP1-96A-GFP
DAPI
50 µm
C
flwFP41 mosaic
RFP(WT) DAPI
50 µm
C'
flwFP41 mosaic
RFP(WT)
50 µm
C''
flwFP41 mosaic
SN
50 µm

Fig S5

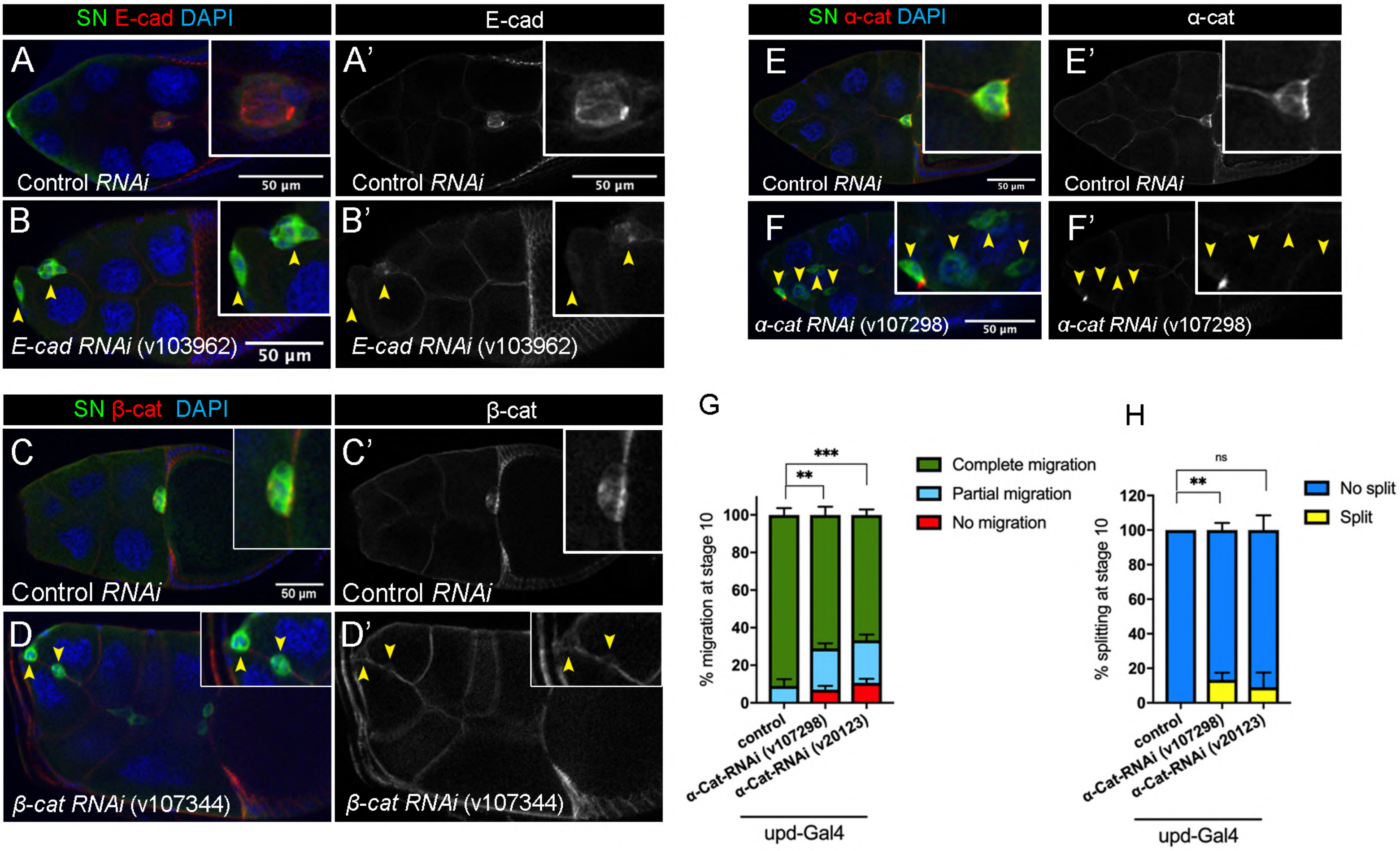

SN E-cad DAPI
E-cad
A
Control RNAi
50 µm
A'
Control RNAi
50 µm
B
E-cad RNAi (v103962)
50 µm
B'
E-cad RNAi (v103962)
SN α-cat DAPI
α-cat
E
Control RNAi
50 µm
E'
Control RNAi
F
α-cat RNAi (v107298)
50 µm
F'
α-cat RNAi (v107298)
SN β-cat DAPI
β-cat
C
Control RNAi
50 µm
C'
Control RNAi
D
β-cat RNAi (v107344)
D'
β-cat RNAi (v107344)
G
***
**
% migration at stage 10
0
20
40
60
80
100
control
α-Cat-RNAi (v107298)
α-Cat-RNAi (v20123)
upd-Gal4
Complete migration
Partial migration
No migration
H
ns
**
% splitting at stage 10
0
20
40
60
80
100
120
control
α-Cat-RNAi (v107298)
α-Cat-RNAi (v20123)
upd-Gal4
No split
Split

Fig S6

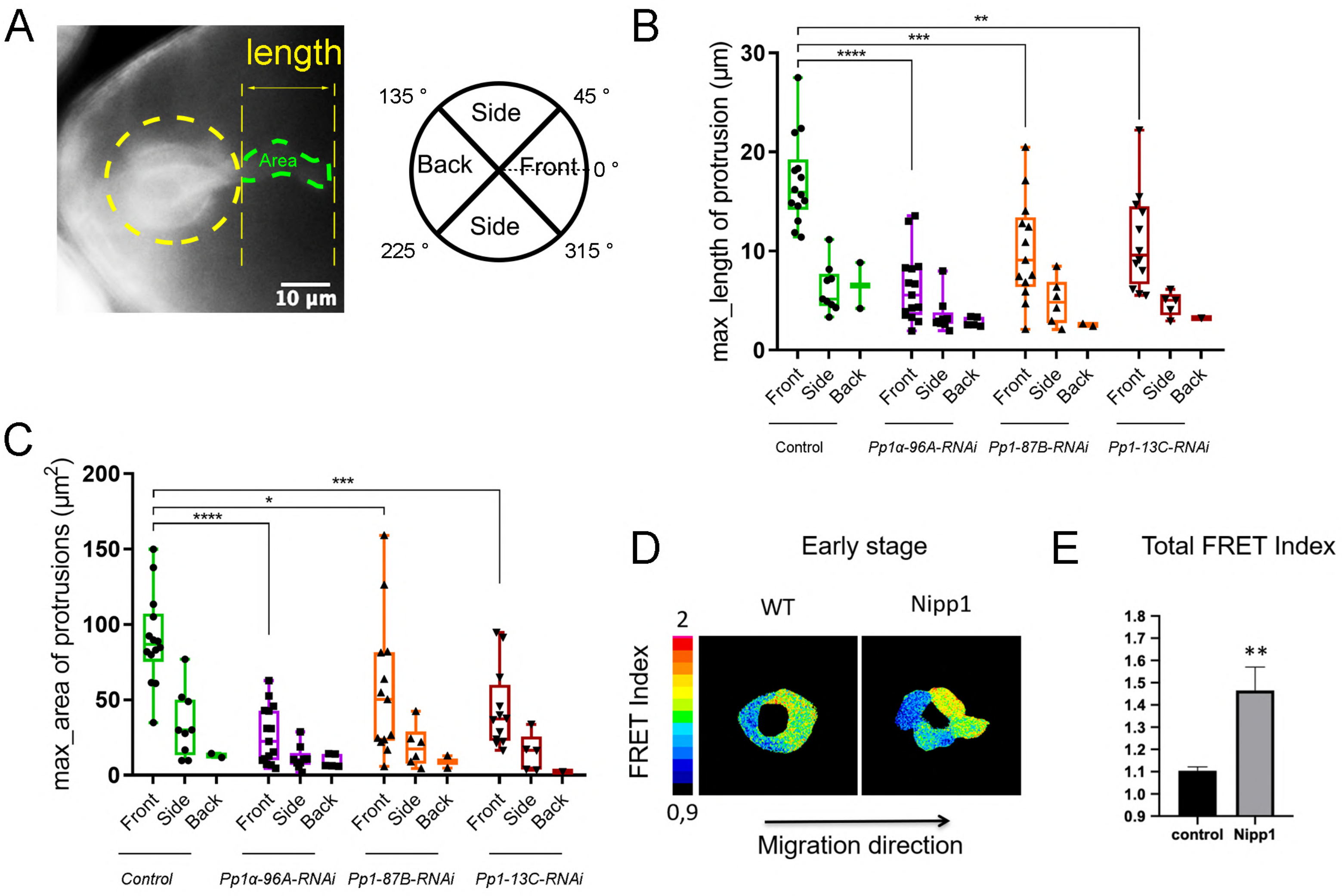

A
length
Area
10 µm
135 °
45 °
Side
Back
Front
0 °
Side
225 °
315 °
B
max_length of protrusion (µm)
30
20
10
0
**
***
****
Front
Side
Back
Control
Pp1α-96A-RNAi
Pp1-87B-RNAi
Pp1-13C-RNAi
C
max_area of protrusions (µm2)
200
150
100
50
0
***
*
****
Front
Side
Back
Control
Pp1α-96A-RNAi
Pp1-87B-RNAi
Pp1-13C-RNAi
D
Early stage
WT
Nipp1
FRET Index
2
0,9
Migration direction
E
Total FRET Index
1.8
1.7
1.6
1.5
1.4
1.3
1.2
1.1
1.0
0.9
**
control
Nipp1

Fig S7

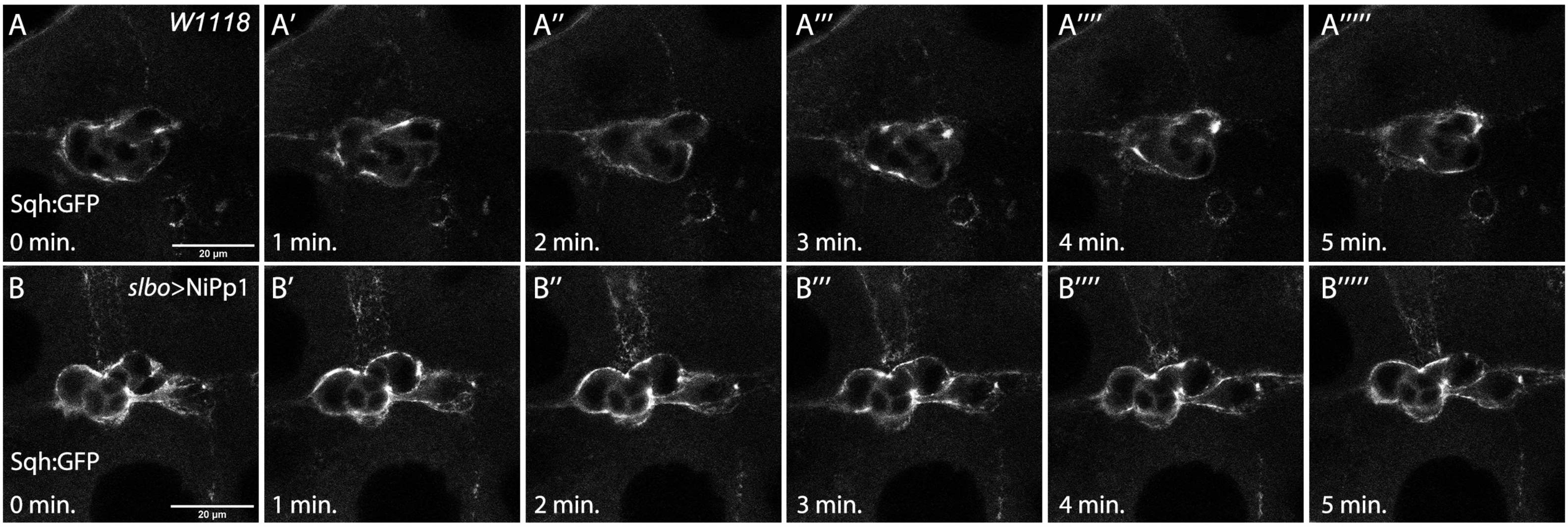
A
W1118
Sqh:GFP
0 min.
20 µm
A′
1 min.
A″
2 min.
A‴
3 min.
A''''
4 min.
A'''''
5 min.
B
slbo>NiPp1
Sqh:GFP
0 min.
20 µm
B′
1 min.
B″
2 min.
B‴
3 min.
B''''
4 min.
B'''''
5 min.

Fig S8

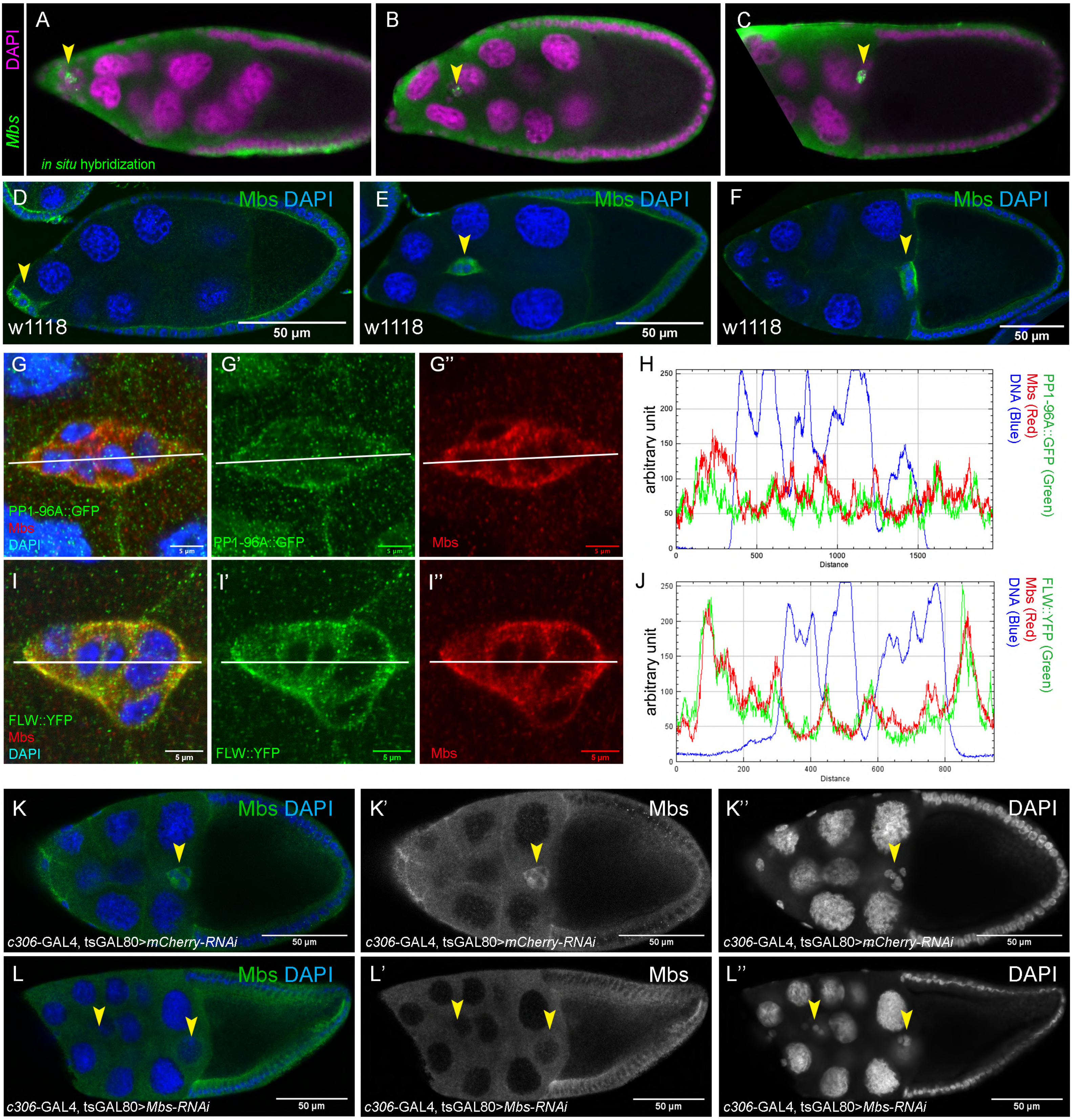

A
B
C
DAPI
Mbs
in situ hybridization
D
E
F
Mbs DAPI
w1118
50 µm
G
G'
G''
PP1-96A::GFP
Mbs
DAPI
5 µm
H
arbitrary unit
Distance
PP1-96A::GFP (Green)
Mbs (Red)
DNA (Blue)
I
I'
I''
FLW::YFP
J
FLW::YFP (Green)
K
K'
K''
c306-GAL4, tsGAL80>mCherry-RNAi
L
L'
L''
c306-GAL4, tsGAL80>Mbs-RNAi

Fig S9

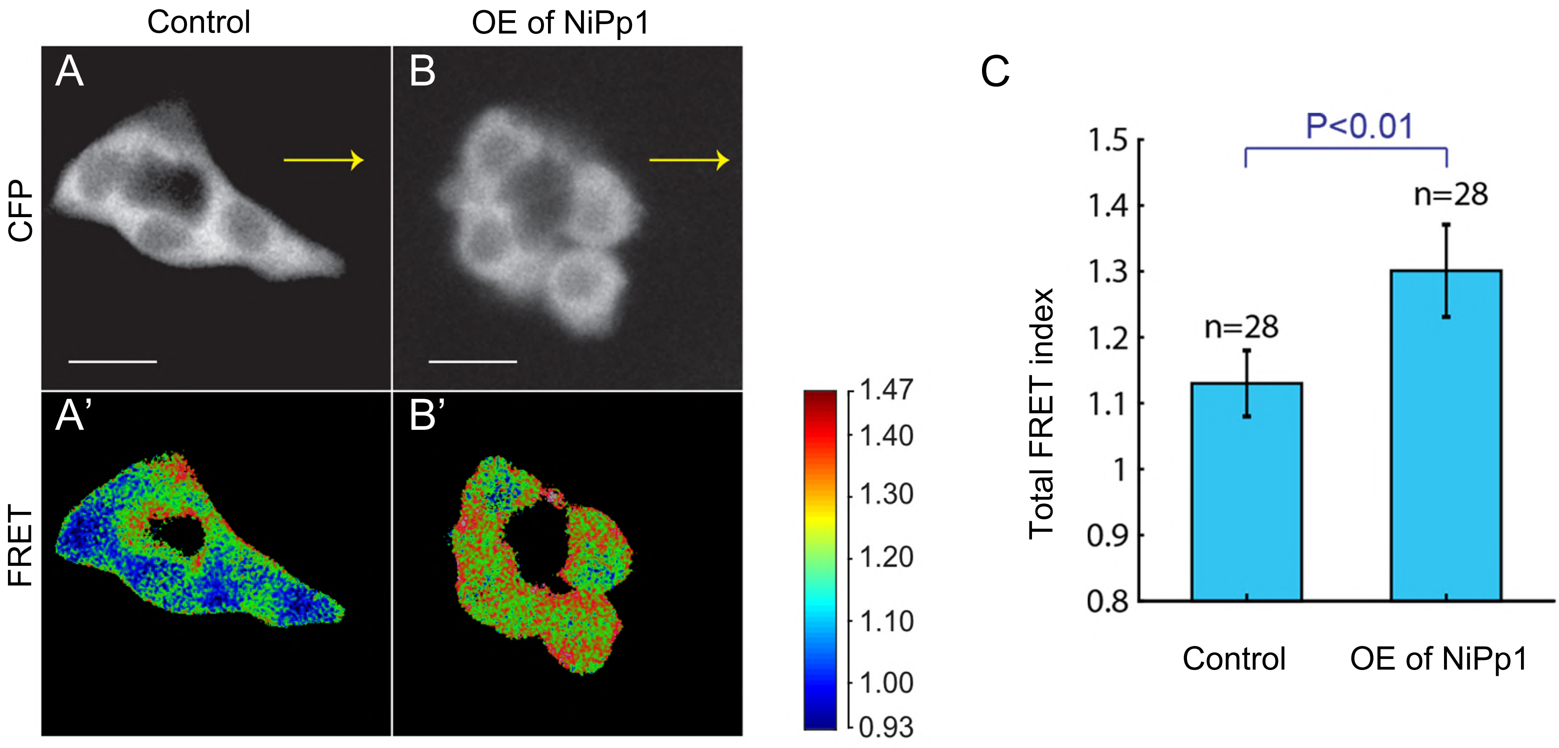

Control
OE of NiPp1
A
B
A'
B'
CFP
FRET
1.47
1.40
1.30
1.20
1.10
1.00
0.93
C
P<0.01
n=28
n=28
Total FRET index
1.5
1.4
1.3
1.2
1.1
1
0.9
0.8
Control
OE of NiPp1
